# Imaging Data-based Model Description Combining Optimal Transport and Phase-field Model

**DOI:** 10.64898/2025.11.30.691255

**Authors:** Tsubasa Sukekawa, Toshiaki Yachimura, Sungrim Seirin-Lee

**Affiliations:** Institute for the Advanced Study of Human Biology(ASHBi), Kyoto University Institute for Advanced Study(KUIAS), Kyoto University, Kyoto 606-8315, Japan; Mathematical Science Center for Co-creative Society, Tohoku University, Sendai 980-0845, Japan; Department of Mathematical Medicine, Graduate School of Medicine, Kyoto University, Kyoto 606-8315, Japan

**Keywords:** Optimal transport, Phase-field model, Cell geometry, Imaging data-driven modeling

## Abstract

The geometrical properties of a cell are not merely passive consequences of cellular function but actively regulate key biological processes during development, morphogenesis, and disease. Although modern live-imaging techniques now allow detailed monitoring of cell morphology, incorporating such complex geometrical information into mathematical models has remained a major challenge. Conventional modeling approaches often rely on artificial cell shape assumptions or purely *in silico* constructions, and discrete imaging data remain fundamentally mismatched with continuous biochemical models based on differential equations. As a result, current models struggle to accurately reproduce biochemical dynamics within realistic, dynamically changing cell geometries. To overcome these limitations, we establish a novel mathematical framework, Imaging Data-based Model Description (IDMD), which integrates Optimal Transport (OT) theory with phase-field (PF) modeling to bridge imaging data and mathematical models. Using live-imaging data from *in vitro* cultured cells and the one-cell *C. elegans* embryo, we demonstrate the versatility of our framework. By directly incorporating real cell geometry into mathematical modeling, our framework provides a powerful new avenue for investigating how geometrical constraints regulate biochemical pattern formation and cell-fate decisions. More broadly, this study highlights a promising direction for integrating modern data science techniques with mathematical modeling, opening new conceptual and methodological possibilities for understanding geometry-driven biological processes in development and disease.

## 1 Introduction

Understanding how the geometrical features of cells contribute to fundamental biological processes is essential for uncovering mechanisms underlying early development and disease. Cell shape is inherently closely linked to function, and changes in geometry are not merely passive consequences but active determinants of cellular behavior, such as differentiation, migration, and intercellular communication (Lecuit and Lenne 2007; Mammoto and Ingber 2010; Seirin-Lee and Kimura 2025). For example, during embryogenesis, spatially coordinated deformations of cells contribute to large-scale tissue patterning and organ formation (Heisenberg and Bellaiche 2013). Similarly, aberrant cell shapes are associated with pathological conditions such as cancer or fibrosis (Friedl and Gilmour 2009), highlighting the importance of quantitatively understanding cellular geometry in both physiological and disease contexts.

Despite advances in live imaging technologies capable of capturing cell morphology and quantitatively analyzing its geometric features (Weigert and et al. 2018), incorporating such complex geometrical information into mathematical models has remained a significant challenge. Conventional mathematical approaches for understanding the influence of cell geometry are often based on artificial cell shape assumptions or entirely *in silico* modeling frameworks. Moreover, conventional modeling methods often rely on overly simplified assumptions and struggle to capture the temporal continuity of actual shape changes in cells. On the other hand, our previous work demonstrated that deviations from biological reality can lead to overlooking critical geometrical factors that regulate cell dynamics (Seirin-Lee et al. 2022).

In general, mathematical modeling faces inherent limitations in simultaneously capturing the spatial and temporal dynamics of actual cell geometry. One major obstacle has been the discrepancy in time scales between experimental imaging data and the *in silico* simulations required for numerical modeling. This mismatch prevents seamless integration of real imaging data into mathematical modeling. To address this gap, a novel mathematical framework is needed that can provide temporally coherent and geometrically meaningful cell shapes. Therefore, in this study, we suggest a new framework that combines Optimal Transport (OT) theory and Phase-field (PF) modeling method, two fundamentally different mathematical tools whose integration has not been established before.

OT is a mathematical framework originally developed to determine the optimal matching between probability distributions (Villani 2009; Santambrogio 2015; Villani 2021). In recent years, OT has gained significant attention in the life sciences, especially for shape matching, image registration, and single-cell omics analysis (Nitzan et al. 2019; Schiebinger et al. 2019; Yang et al. 2020; Moriel et al. 2021; Klein et al. 2023; Yachimura et al. 2024). OT provides interpolation between data distributions, including cell shapes represented as point clouds or density fields. However, the cell geometries described by OT cannot be applied directly to biological problems that are governed by biochemical dynamics. This is because OT-based interpolation operates on discrete data sets, and the resulting geometric information cannot be directly integrated with continuous spatiotemporal biochemical dynamics, which are typically described by systems of differential equations.

We thus combine OT with Phase-field (PF) modeling, known for simulating interface dynamics that provide smooth and continuous descriptions of shape evolution while preserving key physical characteristics (Shao et al. 2010; Nonomura 2012; Seirin-Lee et al. 2016; Akiyama et al. 2018; Seirin-Lee et al. 2020b; Kuang et al. 2022; Seirin-Lee et al. 2022). By combining these two fundamentally different mathematical tools, we establish a method providing temporally precise and biologically validated cell shapes that faithfully reflect the dynamic morphologies observed in living cells. We term this framework Imaging Data-based Model Description (IDMD). It enables the direct incorporation of actual cell geometry, both spatial and temporal, into biochemical models that are typically formulated with differential equations. Our integrated approach has strong potential to significantly advance our understanding of the geometrical principles governing morphogenetic processes, enhance the interpretation of time-lapse imaging data, and facilitate the development of more accurate and mechanistically-grounded mathematical models for developmental and cell biology.

## 2 Materials and Method

### 2.1 Experimental images and segmentation

The experimental images of the *C. elegans* embryo and HL-60 cell were used for testing the IDMD tool. All original *C. elegans* images have the size (*N*_*x*_, *N*_*y*_) = (768, 512), where *N*_*x*_ and *N*_*y*_ denote the numbers of pixels in the horizontal and vertical directions, respectively. All HL-60 images have the same size (*N*_*x*_, *N*_*y*_) = (238, 292). To extract cell shapes as binary images, image segmentation was applied. The Segment Anything Model (SAM) (Kirillov et al. 2023) was used for segmentation.

### 2.2 Numerical algorithm and parameter setting

We solved the phase-field model using a standard explicit finite-difference method implemented in Python. The accuracy of the algorithm was validated using an independently developed code.

The parameters for the phase-field model were chosen such that the simulation time scale technically depends on the extent of cell deformation or movement between images. All details are provided in Appendix H.

## 3 Results

### 3.1 Concept and Workflow of Imaging Data-based Model Description (IDMD)

We briefly introduce the concept and workflow of Imaging Data-Based Model Description (IDMD) (Fig. 1). IDMD enables the simulation of biochemical dynamics within a domain that reflects actual cell deformation based on live imaging data. In particular, we assume a scenario where the temporal snapshot data from live imaging are relatively sparse. In the first stage (Stage I), we transform imaging data into a set of point clouds and interpolate the intermediate shapes of the cells using optimal transport (OT) theory. Then, to ensure data continuity in the second stage (Stage II), we capture the point cloud representation of the cell using a phase-field function, allowing us to obtain a continuous geometric representation of the cell shape. This interpolated data is then integrated with the phase-field model to ensure deformation continuity, thereby representing the spatiotemporal dynamics of an actual cell. In the final stage (Stage III), we demonstrate the versatility of our tool by applying our framework to a biological context. Since our method transforms discrete imaging data into a continuous functional format, it serves as a powerful new modeling tool for investigating the effects of cell geometry on biochemical dynamics in processes such as pattern formation. It is noteworthy to mention that although we use two-dimensional imaging data as validation for our model, the proposed method, in principle, can be applied to data in multiple arbitrary dimensions.

**Figure 1:**
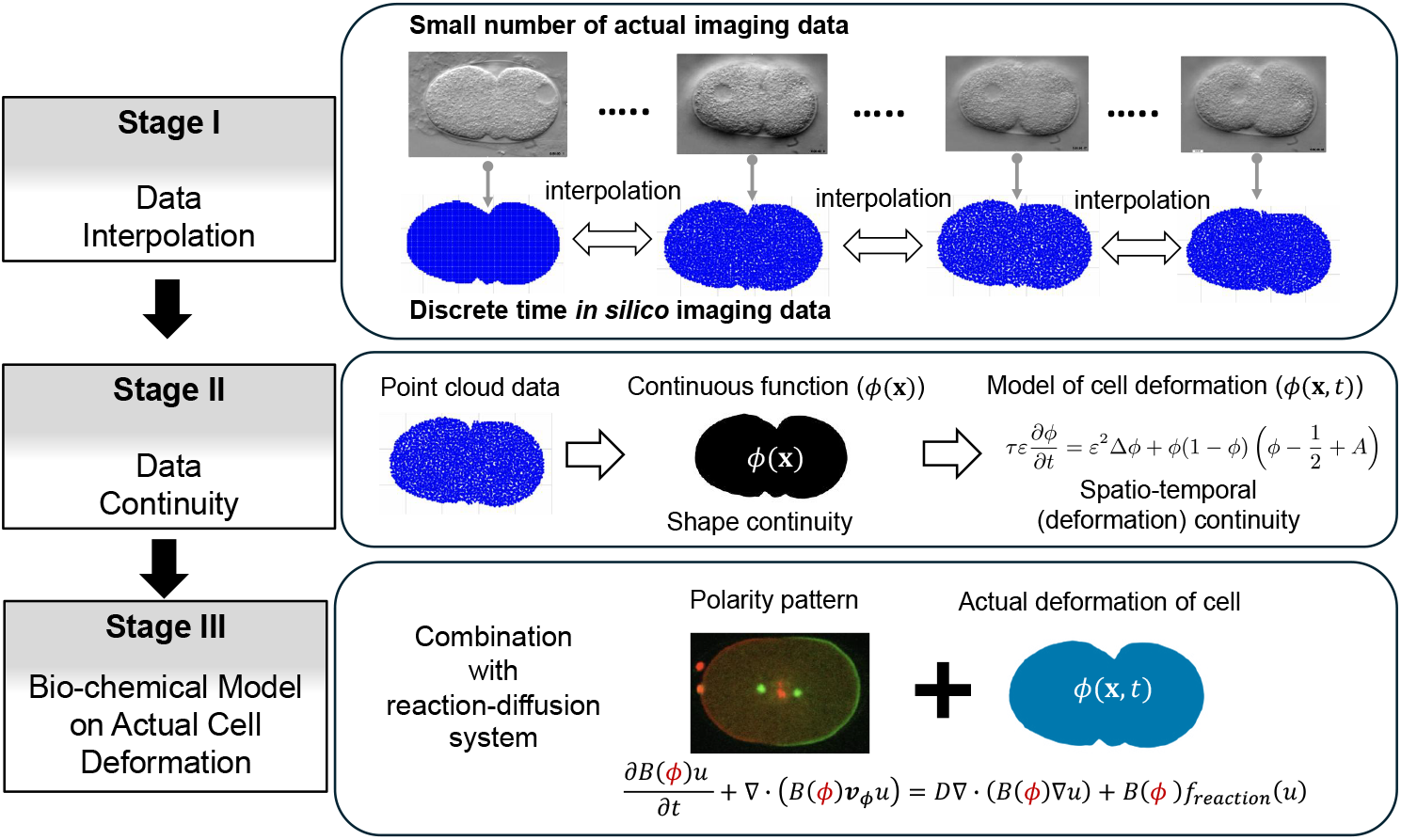
Imaging Data-based Model Description. Stage I: Intermediate cell shapes between imaging time points are interpolated using Optimal Transport (OT) theory. Stage II: Spatiotemporal continuity of cell deformation is reconstructed. Stage III: The framework is integrated with a spatiotemporal biochemical model.

### 3.2 Stage I: Data interpolation of cell shapes using OT from discrete live imaging data

To prepare the imaging data for optimal transport computation, we first segment the cell regions in each image and extract representative points that describe the cell shape. We then model the cell shapes at two time points, *t*_*s*_ and *t*_*g*_, as empirical probability measures *µ*_*s*_ and *µ*_*g*_ in ℝ^2^, corresponding to the source and target distributions, respectively.

To interpolate the shapes between the two time points, we solve the Kantorovich formulation of the optimal transport problem:

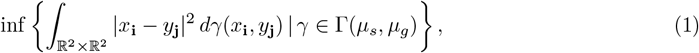

where *µ*_*s*_ and *µ*_*g*_ are empirical measures supported on the observed point sets {*x*_**i**_} and {*y*_**j**_}, respectively. The set Γ(*µ*_*s*_, *µ*_*g*_) denotes all couplings (transport plans) between these measures, and *γ*(*x*_**i**_, *y*_**j**_) represents the amount of mass transported from *x*_**i**_ ∈ supp(*µ*_*s*_) to *y*_**j**_ ∈ supp(*µ*_*g*_). Here, **i** and **j** index the points corresponding to the segmented cell regions at time *t*_*s*_ and *t*_*g*_. When *µ*_*s*_ is absolutely continuous with respect to the Lebesgue measure, Brenier’s theorem (Brenier 1991) guarantees the existence of a unique map *T* that solves the following Monge’s problem:

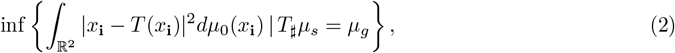

where *T*_*♯*_*µ*_*s*_ denotes the pushforward of *µ*_*s*_ through *T*, defined as *T*_*♯*_*µ*_*s*_(*A*) = *µ*_*s*_(*T*^−1^(*A*)) for measurable sets *A* ⊂ ℝ^2^. Furthermore, the solution *γ* of (1) is given by *γ* = (Id × *T*)_*♯*_*µ*_*s*_. This map *T*, which solves Monge’s problem (2), is referred to as the optimal transport map.

Using the optimal transport map *T*, the interpolation between the source and target distributions can be defined by the formula

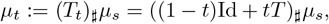

which is known as displacement interpolation or McCann interpolation (McCann 1997) (see Fig. 2A). This interpolation generates a family of intermediate distributions *µ*_*t*_ that smoothly transitions from *µ*_*s*_ to *µ*_*g*_ as *t* varies from 0 to 1.

**Figure 2:**
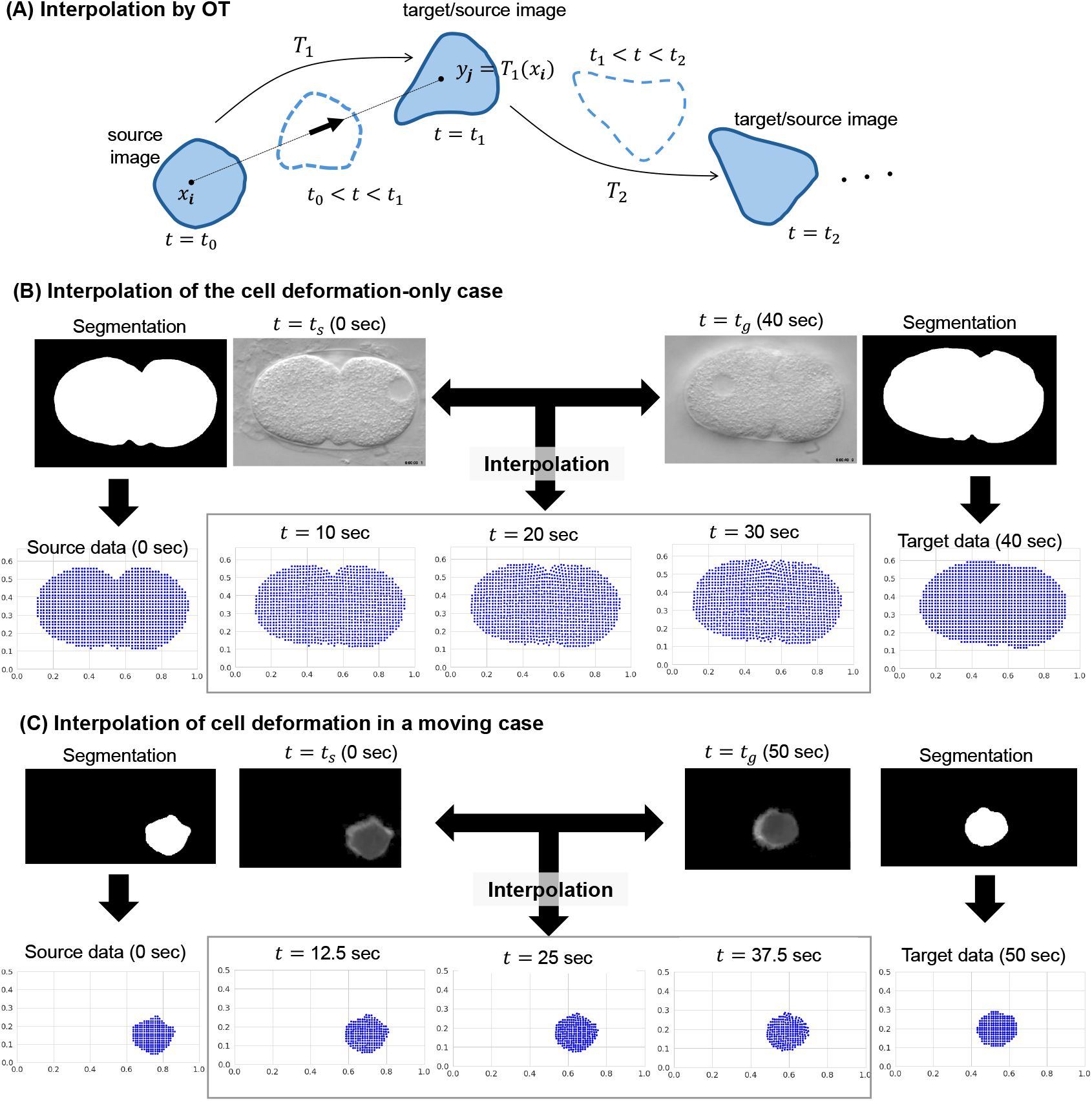
Stage I: Interpolation of cell deformation and movement using optimal transport. (A) Schematic diagram for the interpolation using optimal transport (OT). (B) Interpolation example of the cell deformation-only case for *C. elegans* embryo cell. (C) Interpolation example of the deformation-with-movement case for the HL-60 human cell line.

In the discrete setting, however, an optimal transport map *T* in the Monge sense may not exist, since mass from a single point in *µ*_*s*_ can be split across multiple targets in *µ*_*g*_. To handle this, we employ the barycentric projection map, which defines an approximate transport map based on the optimal coupling *γ* ∈ Γ(*µ*_*s*_, *µ*_*g*_). For each point *x*_**i**_ in the support of *µ*_*s*_, the barycentric projection map is given by

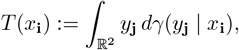

where *γ*(*y*_**j**_ | *x*_**i**_) is the conditional distribution of *y*_**j**_ given *x*_**i**_. This construction provides a well-defined map even in discrete scenarios and allows us to compute the approximate interpolated distributions *µ*_*t*_ in a consistent manner.

Based on the interpolation map *T*_*t*_ = (1 − *t*)Id + *tT*, we define the corresponding velocity field as the time derivative:

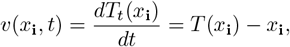

which is constant in *t*. This vector *v*_**i**_ = *T* (*x*_**i**_) − *x*_**i**_ represents the displacement from each point *x*_**i**_ in the source distribution to its corresponding barycentric projection *T* (*x*_**i**_) in the target. The velocity field thus characterizes the direction and magnitude of deformation at each point, and serves as the basis for constructing intermediate cell shapes between *µ*_*s*_ and *µ*_*g*_. We refer to supporting information about the numerical algorithms related to the velocity field.

Using the velocity vectors *v*_**i**_, we reconstruct intermediate cell shapes by linearly interpolating each point along its displacement path. To evaluate the effectiveness of this approach, we tested two representative cases using live imaging data: a deformation-only case involving *C. elegans* embryo, and a deformation-with-movement case involving the HL-60 human leukemia cell line (see Figs. 2B, C). These examples demonstrate that optimal transport interpolation can capture not only smooth deformations of cell shape but also translational movements across time points.

When multiple time points *t*_0_, *t*_1_, …, *t*_*N*_ are available, we represent the corresponding cell shapes as a sequence of empirical measures 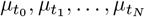. For each consecutive pair 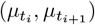, we solve the optimal transport problem (1) to obtain the corresponding barycentric projection map *T*_*i*_, and define the velocity field 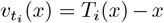, which describes the optimal displacement from time *t*_*i*_ to *t*_*i*+1_. This yields a sequence of velocity fields 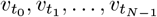, each representing shape changes over its respective time interval. To construct a time-continuous description, we define a piecewise velocity field 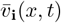 by specifying vector fields 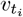 on each interval [*t*_*i*_, *t*_*i*+1_].

### 3.3 Stage II: Data continuity by phase-field model

While the velocity fields obtained from optimal transport provide information about cell deformation over time, the interpolated cell data remain in the form of discrete points. They cannot be directly incorporated into mathematical models formulated with partial differential equations. To bridge this gap, we introduce a method to convert the discrete geometric data into a smooth, continuous representation of cell shape over both space and time. Our approach uses a phase-field model, which enables the representation of cell geometry as a continuous scalar field and provides a natural way to simulate morphological evolution.

The core concept is to interpret the point-wise deformation of the cell, represented by the velocity field from optimal transport, as a traveling wave propagating through a continuous medium (Fig. 3A). With this perspective, each point in the velocity field corresponds to a local interface movement that can be modeled by the speed of a traveling wave. Below, we provide an intuitive summary of the mathematical framework that enables this translation from discrete motion vectors to continuous field evolution.

**Figure 3:**
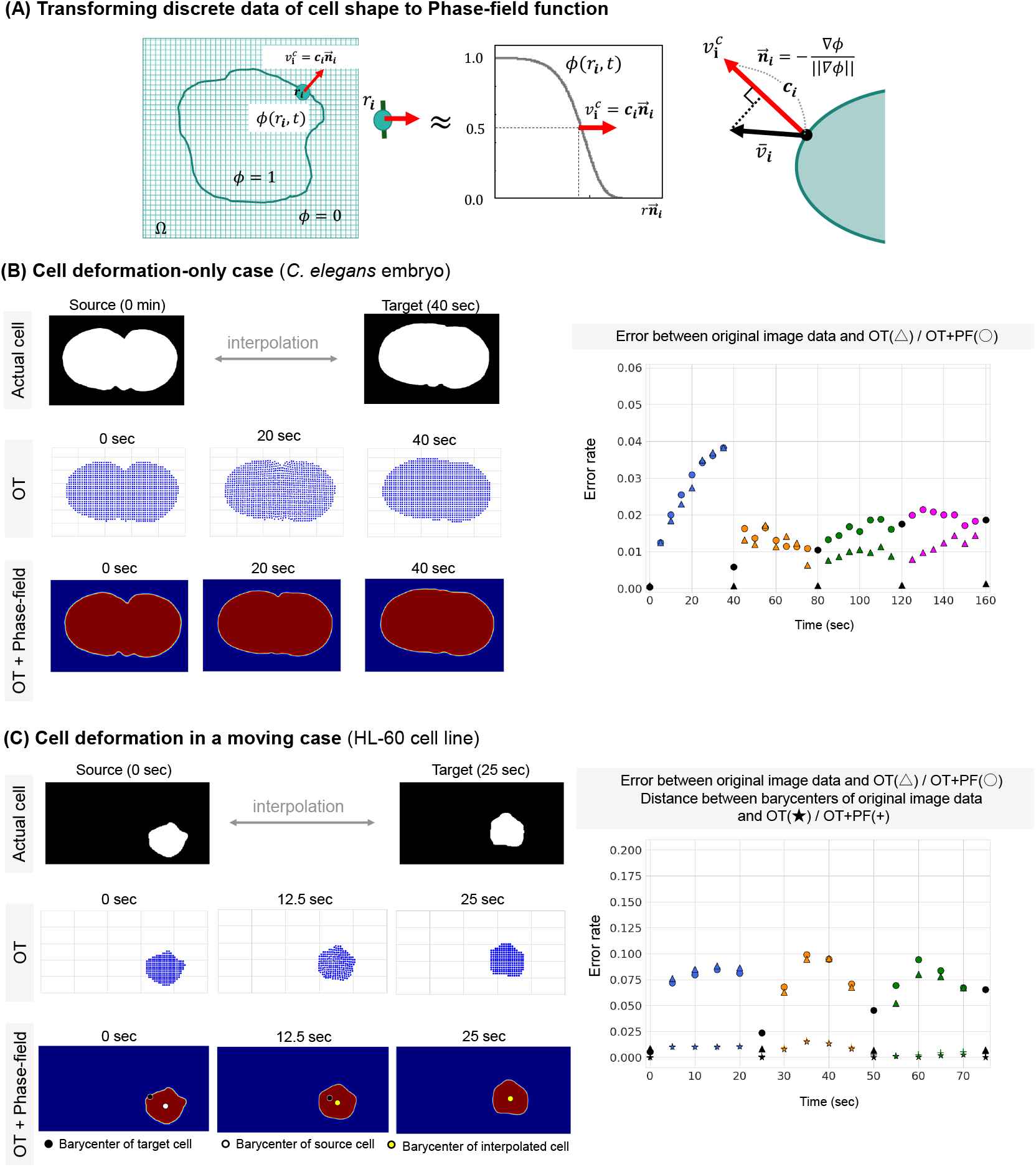
Stage II: Data continuity by phase-field model. (A) Core idea connecting interpolation data obtained by optimal transport(OT) with the phase-field model(PF). (B-C) Results of deformation-only case using *C. elegans* embryo and deformation in a moving case using the HL-60 cell line. The left panels show the error rate between the original image data and the output of our method. The error function, (*f, g*), is given in the equation (6). The error of the barycenter location was calculated by the Euclidean distance.

To model the continuous deformation of the cell shape, we employ a phase-field equation that governs the temporal evolution of a scalar field *ϕ*(*x, t*), which smoothly transitions between 0 (outside the cell) and 1 (inside the cell). The governing equation is given by (see Seirin-Lee et al. (2022) in more detail):

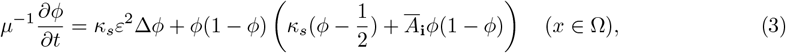

where Ω = [0, *L*_*x*_] × [0, *L*_*y*_] is a domain of the system and image size, *µ* is the mobility constant of interface motion describing cell shape, and *κ*_*s*_ is the surface tension coefficient. Since *ε >* 0 is a parameter determining the width of the interface, we take it to be a sufficiently small value. A schematic image of the phase-field model of cell deformation is given in Fig. 3A. This equation describes the motion of the cell boundary as an evolving interface in a bistable medium, with the term involving *Ā*_**i**_ that modulates the direction and speed of the deformation at each point.

In the case of an effectively infinite domain, the equation supports traveling wave solutions. The speed 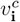 of such a traveling wave is given by

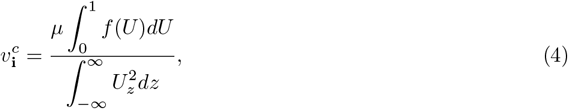

where *U* (*z*) = *ϕ*(*r*_**i**_, *t*) with 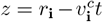 and *f* (*U*) = *U* (1 − *U*) *κ*_*s*_(*U* − 1*/*2) + *Ā*_**i**_*U* (1 − *U*) (see Appendix A for more details). Based on the equation (4), the coefficient *Ā*_**i**_ can be explicitly linked to the traveling wave speed 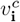 by

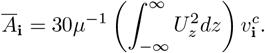

We carry out the model modification and rescaling of the original model (3) to avoid calculating the uncertain quantity included in *Ā*_**i**_. With denoting 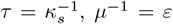, and replacing *A*_**i**_(*x, t*) with 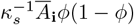, we obtain

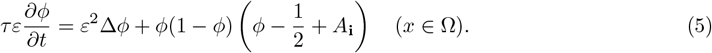

Note that we can obtain the exact form of the traveling wave solution to (5), which allows us to determine the parameter *A*_**i**_ from the wave speed. We define *A*_**i**_ as follows:

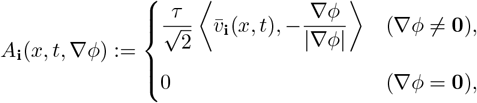

where 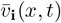 is a piecewise vector field that appeared in the previous section. Namely, this velocity field corresponds to the actual cell geometry obtained from the cell live image data. This velocity field 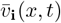 enables us to capture the piecewise continuous evolution of cell shape across the time sequence. ⟨·, ·⟩ denotes the inner product on ℝ^2^, and **0** is the zero vector. This formulation ensures that *A*_**i**_ is proportional to the projection of 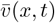 onto the outward unit normal vector −∇*ϕ/*|∇*ϕ*|, directing the interface motion according to 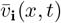 (Fig. 3(A) right panel). By properly choosing the parameter *τ* (the reciprocal of surface tension), we can satisfy the bistability condition and fine-tune the cell shape with minimal error caused by the surface tension effect in the phase-field model. This approach provides a comprehensive framework for modeling continuous changes in cell shape across a series of images by integrating information from all time points, providing a smooth cell shape.

Next, we confirm our method for the deformation-only case and cell deformation in a moving case. Figs. 3B and C show the corresponding results for a *C. elegans* embryo and an HL-60 human cell line, respectively. To evaluate how accurately our method captures the original cell image data, we defined the following error function:

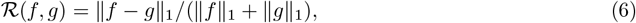

where ∥ · ∥_1_ is the norm of *L*^1^(Ω), *f* and *g* indicate the original image data and OT/OT+PF data, respectively.

We first compared the error rates between the cell image data and the OT+PF case. In both the deformation-only and deformation-with-movement settings, the error rate was less than 10% (Figs. 3B and C, left panels). In the case of deformation with movement, however, relatively higher errors were observed compared to the deformation-only case. These errors are likely synergistically amplified by inaccuracies in cell positioning, but they can be reduced by selecting shorter time-step data between the source and target cells.

In addition, the OT+PF model did not show notable differences from the interpolation results obtained with OT alone. This indicates that our OT+PF framework not only maintains a comparable level of accuracy to OT but also effectively converts the inherently discrete imaging data into a continuous representation. This demonstrates that our integrated OT+PF model provides a seamless bridge between discrete imaging data and continuous geometric representations.

### 3.4 Stage III: Modeling of bio-chemical/cellular dynamics on actual cell deformation

Next, to demonstrate the versatility of our tool, we applied it to modeling in a biological context. Although our model can be applied to various biological situations, its strength is the capacity to combine *in silico* reaction-diffusion models describing biochemical/cellular dynamics onto actual cell/organ geometries, including deformation. This is because our method provides a direct transformation of cell geometry into a phase-field function, which can be further extended directly by incorporating the phase-field function into biochemical/cellular dynamics reaction-diffusion models.

The mathematical model combining a phase-field model and reaction-diffusion system has been extensively used for understanding single-cell dynamics (Aras et al. 2017; Saito and Sawai 2021; Imoto et al. 2021; Seirin-Lee 2021; Wang et al. 2017) and multi-cellular or tissue dynamics (Seirin-Lee 2016). The basic mathematical format is given by

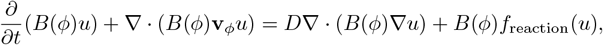

where *u*(**x**, *t*) is the concentration of a substrate and *D* is its diffusion coefficient. The function *B*(*ϕ*) defines the substrate’s localization: for a substrate in cytosol, *B*(*ϕ*) = *ϕ*, and for a substrate in the membrane, *B*(*ϕ*) = *αϕ*(1 − *ϕ*), where *α >* 0 is a constant.

The advection term, ∇·(*B*(*ϕ*)**v**_*ϕ*_*u*), describes the transport of the substrate driven by cell deformation. A crucial advantage of our framework is that the velocity vector **v**_*ϕ*_ is not a theoretical assumption but is directly calculated from the velocity fields 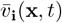 interpolated by OT in Stage I. This direct linkage ensures that the influence of the cell’s observed motion is precisely incorporated into the simulation, naturally compensating for complex interface dynamics. Furthermore, the approach is highly versatile, as the governing equation holds for any reaction term *f*_reaction_(*u*).

To demonstrate the capability of this framework, we simulated the PAR-2 polarity mathematical model, previously studied in Seirin-Lee and Shibata (2015); Seirin-Lee et al. (2020a), on the actual deforming geometry of a one-cell-stage *C. elegans embryo* (Fig.4). This example illustrates that polarity of the cell membrane is formed in the convex regions rather than the concave regions, and that the maximal PAR-2 concentration level is influenced by cell deformation (The dotted circles in Fig.4). These results indicate that dynamic deformation of the cell may critically affect robust polarity formation and serve as an important factor regulating cell fate.

## 4 Discussion

Although geometric constraints of a cell play a critical role in determining cell fate, experimental approaches alone remain insufficient to fully understand how geometry is precisely related to the overall process of fate determination. Consequently, a combination of experimental and theoretical approaches is important. Nevertheless, conventional theoretical models have typically relied on artificially defined geometries, such as perfect spheres or ellipsoids, failing to capture the complexity of actual cell shapes. In this study, we have developed a novel mathematical framework called Imaging Data-based Model Description (IDMD) that integrates optimal transport (OT) theory with the phase-field (PF) method. This enables us to incorporate actual cell shapes from live imaging data. This framework makes it readily feasible to simulate systems of differential equations on realistic, dynamic cell domains. The core of our framework lies in the synergistic combination of OT and PF. OT provides a powerful method for temporal interpolation between discrete image frames, allowing for robust simulations even with limited imaging data, thereby reducing experimental and computational costs, data size, and adverse phototoxic effects from laser excitation. The key idea is to combine PF modeling with OT, which allows us to smooth spatially discrete data and convert it into a continuous spatial representation. For example, unlike direct OT interpolation that can produce unnatural shapes, our method generates smooth cell morphologies (Fig. S1).

While our framework effectively captures cell deformation, its performance on complex cell migration trajectories indicates an area for future improvement. The current implementation uses McCann’s interpolation (McCann 1997), and its trajectories are constructed by creating a linear path between two adjacent time points in Wasserstein space. While effective for simple deformations, this local approach can produce trajectory errors for highly non-linear movements observed in real data. A straightforward solution is to increase the temporal resolution of the input data, but a more sophisticated approach would be to enhance the interpolation method itself. To this end, exploring non-linear, OT-based interpolation methods presents a promising direction. In contrast to our local method, recent advances in spline interpolation in Wasserstein space utilize the entire sequence of shapes to generate a globally smooth trajectory (Benamou et al. 2019; Chen et al. 2018; Justiniano et al. 2024). By considering the full time-series data at once, these frameworks can capture complex, non-linear dynamics more faithfully. Integrating these advanced global frameworks into our IDMD pipeline could substantially improve the accuracy of migration trajectory predictions and further broaden the applicability of our method.

Finally, we would like to emphasize that our study demonstrates how modern data science techniques can be integrated with mathematical modeling to create a new mathematical framework. The application of mathematical modeling to biology dates back more than 220 years to Malthus’s model of 1798 (Malthus 1798). Nevertheless, conventional mathematical approaches have remained largely confined to classical differential-equation-based frameworks that have changed little over time, and thus have not progressed as rapidly as experimental biology or computer science. Our study provides one of the first examples showing a possible direction for the methodological integration of data science and mathematical modeling.

Our framework, which incorporates actual geometrical information from cell imaging data into mathematical models, may open a new avenue in biology. While geometric constraints have long been recognized as central to cell fate decisions, their mechanistic roles have remained largely unexplored due to experimental limitations. By embedding real cell geometry directly into mathematical models, our framework overcomes these barriers and provides a powerful tool for uncovering how geometry governs pattern dynamics. More importantly, this framework does not merely complement existing biological approaches. It could open a new avenue for addressing fundamental questions that were previously inaccessible. Applying this methodology to concrete biological systems will not only deepen our understanding of cell behavior but may also redefine how geometry-driven mechanisms are studied in modern biology.

## Data availability

All relevant data are included in the manuscript and the appendix information files.

## Code availability

Numerical codes and all other data are available from the authors on reasonable request.

## Author Contributions

SSL and TY initiated, designed, conducted, and supervised the research. TS conducted the research and analyzed the data. TS and SSL drafted the initial version of the manuscript, and SSL was primarily responsible for major revisions. All authors reviewed and revised the final version.

## Acknowledgements

We thank Dr. Takahiro Hiraga for performing the segmentation of the live imaging data. This work was supported by the Japan Society for the Promotion of Science (JSPS) KAKENHI Grant-in-Aid for Transformative Research Areas (A) (22H05110) to SSL, and JST, PRESTO Grant Number JPMJPR24KD, Japan to TY.

## Appendix

### A Calculation of the traveling wave speed in the PF model

Omitting the index **i**, the governing equation of the PF model in Eq. (3) is given by

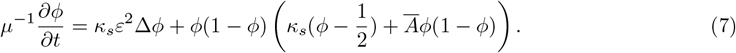

Instituting *U* (*z*) = *ϕ*(*r, t*) with *z* = *r* − *v*^*c*^*t* into the equation (7), we obtain

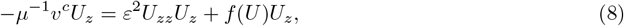

where *f* (*U*) = *U* (1 − *U*)(*U* − 1*/*2 + *Ā U* (1 − *U*)). Then, Eq. (8) can be calculated as follows:

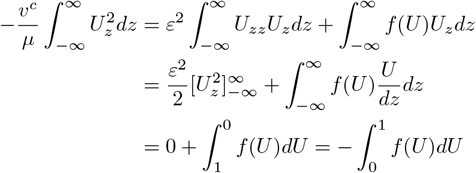

Therefore, we obtain

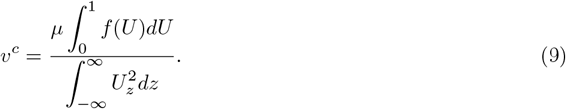

Now, we calculate the integral term:

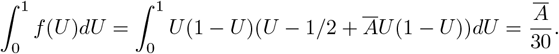

Finally, we obtain

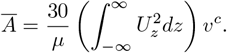

### B Polarity model in *C. elegans* embryo

For the test simulation in Fig. 4, we used the following PAR-2 polarity model (Seirin-Lee and Shibata 2015; Seirin-Lee 2021). We choose the cell membrane periphery model (Morita and Seirin-Lee 2021).

**Figure 4:**
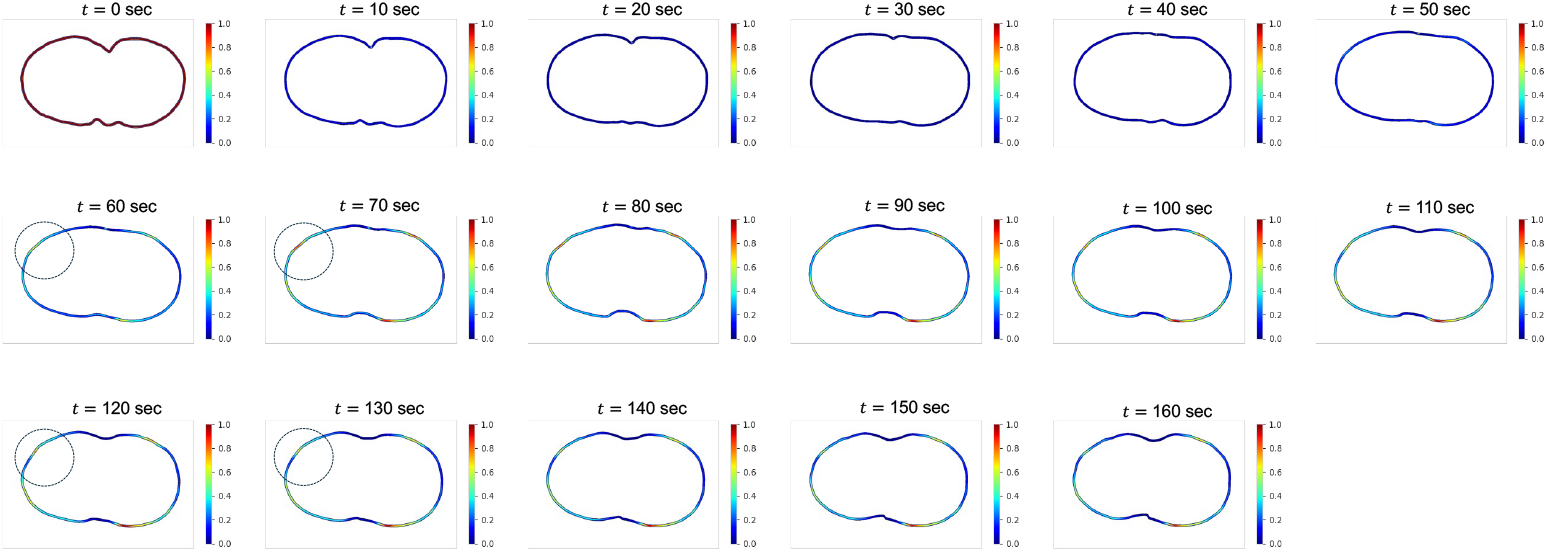
Stage III: Modeling of bio-chemical dynamics on actual cell deformation. Example of the PAR-2 polarity model in a *C*.*elegans* embryo with actual cell deformation. See Appendix B for the detailed model equations. The color bar indicates the concentration of PAR-2. The dotted circles highlight examples showing that the maximum PAR-2 concentration changes depending on cell deformation.

Let us denote the concentrations of PAR-2 in the membrane and cytosol in the cell membrane periphery by *u*(**x**, *t*) and *v*(**x**, *t*), respectively. Then the model equations are given by

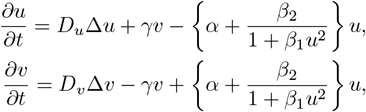

where *D*_*u*_ and *D*_*v*_ are diffusion coefficients of PAR-2 on cell membrane and cytosol, respectively. The final model equations, combined with actual cell deformation data, are given by

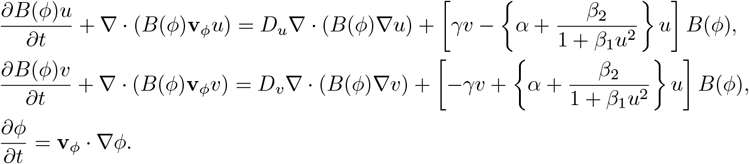

### C Unnatural deformations appearing in OT interpolation

In the optimal transport theory, displacement interpolation (McCann 1997) can produce intermediate cell shapes using two images. However, standard implementations may yield unnatural deformations during interpolation. Figure S1 presents results of displacement interpolation for cell shape data, where we refer to the algorithm in (Solomon et al. 2015). These deformations appear to disregard certain physical properties of the cell membrane.

### D Velocity field associated with OT from imaging data

Hereafter, we outline the computation of the velocity field 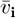 in the main text. We consider binary imaging data in which the cell shape and extracellular area are distinguished in black and white. Let *d* denote the spatial dimension of the data; *d* = 2 and *d* = 3 correspond to pixel and voxel data, respectively. In this section, we mainly focus on the explanation in the case *d* = 2. The data are regarded as discrete probability measures. Let *µ*_*s*_, *µ*_*g*_ be the source and target probability measures associated with the source and target images.

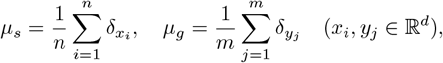

where *x*_*i*_, *y*_*j*_ are support points for *µ*_*s*_ and *µ*_*g*_, respectively, and *δ*_*x*_ is the Dirac measure at position *x*. Also, *n* and *m* are the numbers of support points for the source and target measures, respectively. In practice, *x*_*i*_ and *y*_*j*_ are identified with pixel positions of the images. Let *C* = (*C*_*ij*_) and *P* = (*P*_*ij*_) be real matrices of dimension *n* × *m*. Throughout, we consider *C*_*ij*_ = |*x*_*i*_ − *y*_*j*_|^2^.

**Figure S1:**
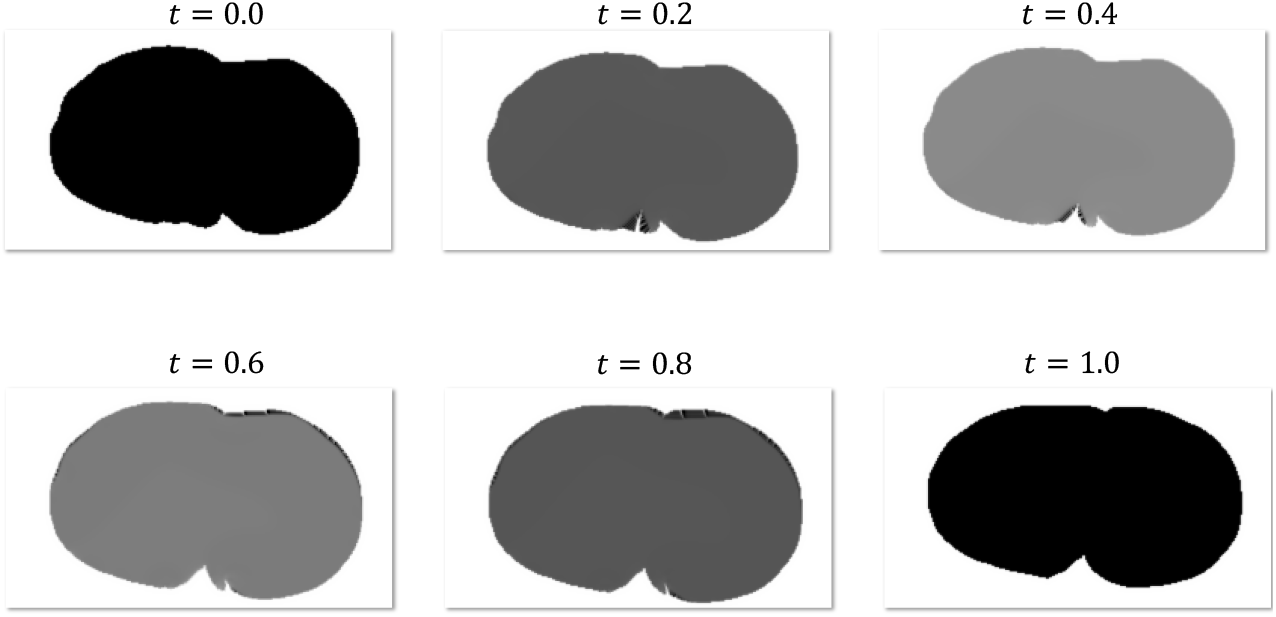
The example of the unnatural deformations observed in the OT interpolation. The top left figure is the source, and the bottom right figure is the target cell shape. The cell shape is represented in grayscale, with values ranging from 0 to 1. At each *t*, the function’s maximum value is normalized to 1 for visualization. A crack-like deformation is observed at *t* = 0.2, and the sharpened structure is observed at *t* = 0.8.

We consider the following minimization problem:

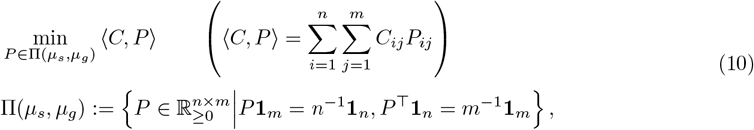

where *P* is regarded as vector with non-negative *n* × *m* elements, **1**_*n*_ ∈ ℝ^*n*^ is vector of ones, and *P*^⊤^ denotes the transpose of *P*. The problem (10) is the Kantorovich relaxation formulation of the optimal transport problem, which is a linear programming problem with variables *P*_*ij*_. *C*_*ij*_ and *P*_*ij*_ are regarded as unit costs and relative amounts of transportation from *x*_*i*_ to *y*_*j*_, respectively. The objective function ⟨*C, P* ⟩ is the total transportation cost from *µ*_*s*_ to *µ*_*g*_. The conditions *P* **1**_*m*_ = *n*^−1^**1**_*n*_, *P*^⊤^**1**_*n*_ = *m*^−1^**1**_*m*_ are referred to as mass conservation constraints. These constraints require that the amount of transportation from *x*_*i*_ must be *n*^−1^, and the amount of transportation to *y*_*j*_ must be *m*^−1^. The feasible set Π(*µ*_*s*_, *µ*_*g*_) is called the transportation polytope. We note that Π(*µ*_*s*_, *µ*_*g*_) is a compact set, and the objective function is continuous, so there exists a minimizer to the problem (10). Let *P*^∗^ denote such a minimizer to the problem (10). *P*^∗^ is called an optimal transport plan. If *P*^∗^ has exactly one positive element in each row, then it induces a map, which is called an optimal transport map.

Our objective is to construct the velocity field by using the optimal transport map. However, due to mass constraints, we cannot obtain the optimal transport map when *n < m*. In this case, *P*^∗^ induces mass splitting, which precludes map construction. Instead of an optimal transport map, we employ the following barycentric projection map:

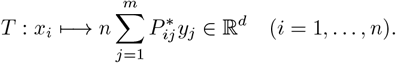

Note that *T* is defined only on {*x*_1_, …, *x*_*n*_} ⊂ ℝ^*d*^. Let *v*_*i*_:= *T* (*x*_*i*_) − *x*_*i*_ ∈ ℝ^*d*^, and define a family of vector-valued measure as follows:

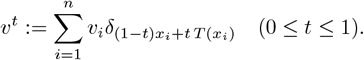

Note that the support of *v*^*t*^ is defined only on points (1 − *t*)*x*_*i*_ + *t T* (*x*_*i*_) (*i* = 1, …, *n*), thus *v*^*t*^ itself cannot be directly combined with the phase-field equation (5). To obtain a vector field defined on the entire domain Ω, we perform nearest-neighbor interpolation on *v*^*t*^.

For three or more images, the above construction extends straightforwardly. Let *µ*_*k*_ (*k* = 0, 1, …, *N*) be probability measures corresponding to *k*-th image, i.e.,

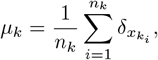

where 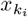 is support point for *µ*_*k*_, and *n*_*k*_ is the number of support points for *µ*_*k*_. For *k* = 1, …, *N*, let *T*_*k*_ be the barycentric projection map obtained from an optimal coupling between *µ*_*k*−1_ and *µ*_*k*_, and denote 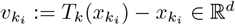. The *k*-th vector-valued measure is defined by

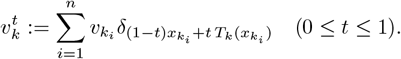

Nearest-neighbor interpolation is applied on 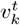 and let *v*_*k*_(*x, t*) denote the resulting vector field. Finally, we obtain the velocity field defined on all intervals:

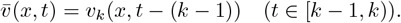

### E Bidirectional method

The accumulation of errors is mitigated by decreasing *ε* or increasing *τ*, thereby reducing the effect of interfacial tension. However, if *ε* is too small, the spatial step width in the numerical simulation must also be reduced, which increases computational cost. Conversely, if *τ* is too large, the bi-stability condition cannot be satisfied, potentially leading to numerical instability.

Instead of modifying the parameters, we propose a bidirectional method, which computes intermediate cell shapes in both the forward and backward directions. For simplicity, consider the two cell shape data labeled No. 0 and No. 1. In the forward direction, the intermediate cell shape is inferred by solving the phase-field model with No. 0 as the source and No. 1 as the target, which is the standard setting. Conversely, in the backward direction, the roles of the datasets are interchanged—No. 0 as the target and No. 1 as the source—and the phase-field model is computed accordingly. Hereafter, the methods are referred to as the “forward method” and the “backward method”, respectively.

Let *ϕ*_f_(*x, t*) and *ϕ*_b_(*x, t*) denote the solutions obtained by the forward and backward methods, respectively. We combine these solutions in analogy to linear interpolation, as follows:

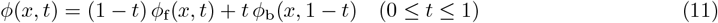

We note that the time directions are opposite in the forward and backward methods. Namely, *ϕ*_f_(*x*, 0), *ϕ*_f_(*x*, 1) correspond to No. 0 and No. 1, whereas *ϕ*_b_(*x*, 0), *ϕ*_b_(*x*, 1) correspond to No. 1 and No. 0, respectively. This is the reason the term *ϕ*_b_(*x*, 1 − *t*) appears in (11). The coefficients (1 − *t*) and *t* are used as the weights of the functions.

We note that the bidirectional method is capable of interpolating between shapes, whereas the forward method does not, in general, provide a rigorous interpolation. This follows from *ϕ*(*x*, 0) = *ϕ*_f_(*x*, 0) and *ϕ*(*x*, 1) = *ϕ*_b_(*x*, 0). The functions *ϕ*_f_(*x*, 0) and *ϕ*_b_(*x*, 0) can completely match to the cell shapes No. 0 and No. 1, respectively. Consequently, *ϕ* can exactly match No. 0 and No. 1 at *t* = 0 and *t* = 1.

Fig. S2 compares the numerical results of the forward method and the bidirectional method. In the bidirectional method, the phase-field function shown in the figure is a linear combination of the forward and backward solutions; consequently, regions with values close to 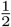 may appear at *t* = 0.5 (Fig. S2). This suggests that the intracellular regions obtained by the two methods overlap in those areas. The cell shape can then be extracted from these results by binarization. We note that the bidirectional method can be extended to three or more images by applying the forward and backward methods to two consecutive images.

In contrast to the forward method, the bidirectional method can perform interpolation. This is a significant property because, in the forward method, the phase-field function does not, in general, exactly match the target cell shape. This is because there is an accumulation of error due to the effect of surface tension. In particular, the magnitude of the error may increase with the number of images. On the other hand, the bidirectional method prevents the accumulation of the error by using the phase field function in the back direction *ϕ*_b_ and the weights *t* and (1 − *t*). Roughly speaking, the error accumulation in the forward method is canceled by the backward method.

**Figure S2:**
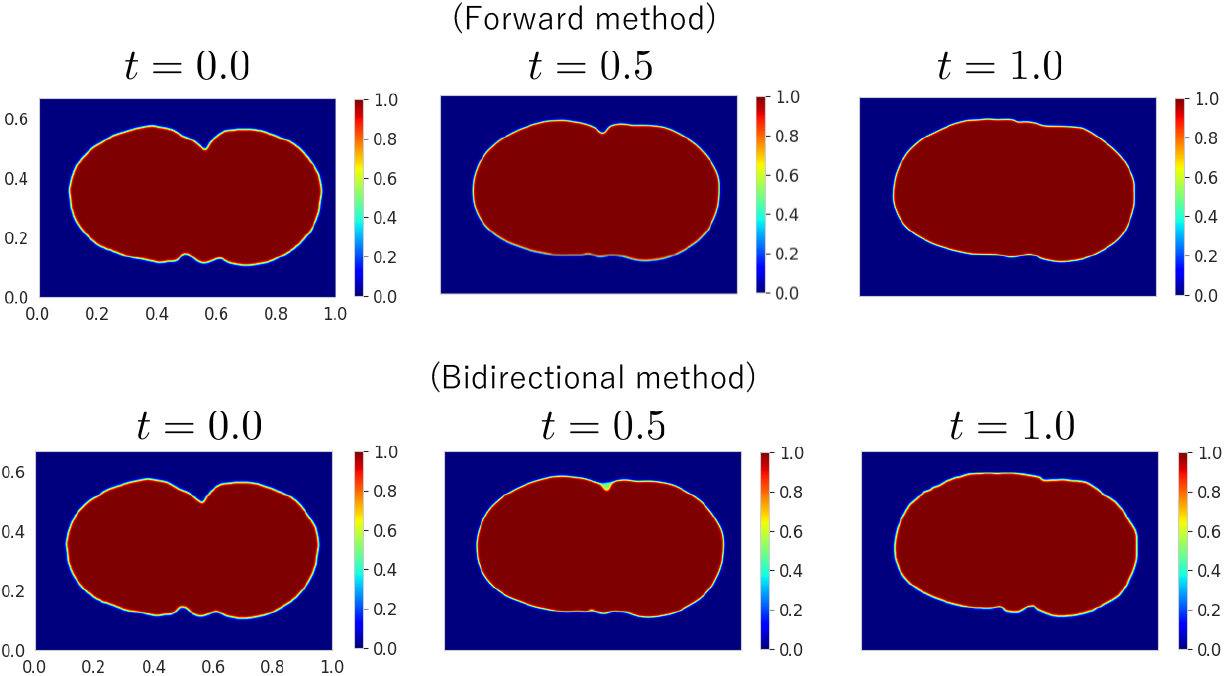
The phase field functions obtained by forward and bidirectional methods.

Fig. S3 presents the numerical results of the forward and bidirectional methods for comparison to the intermediate images between No.0 and No.8 of *C. elegans* (see Fig. 2 in the main paper), where the result of the forward method is the same as the one in Fig. 2 in the main paper. At the final image (No. 8), the error under the bidirectional method is close to zero. We note that the tiny error is induced by applying a Gaussian filter to *ϕ*_b_(*x*, 0), which corresponds to the image No. 8 in this case. By contrast, the error for No. 8 under the forward method is comparatively larger. These observations suggest that the bidirectional method mitigates error accumulation.

### F Extrapolation of cell shapes

In this section, we present results on the extrapolation of HL-60 cell shape data. Extrapolating shapes is, in general, more challenging than inferring intermediate shapes. Since the target shape is unknown in extrapolation, a velocity field defined outside the observed time interval cannot be constructed by the same procedure used for interpolation.

To construct the vector field, we extend the velocity field numerically in a heuristic manner.

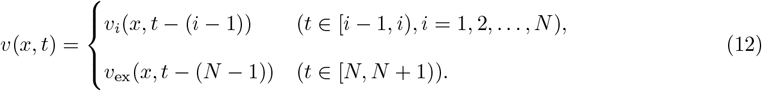

The final vector field *v*_ex_ is defined as follows:

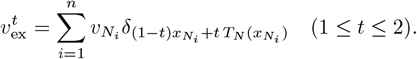

**Figure S3:**
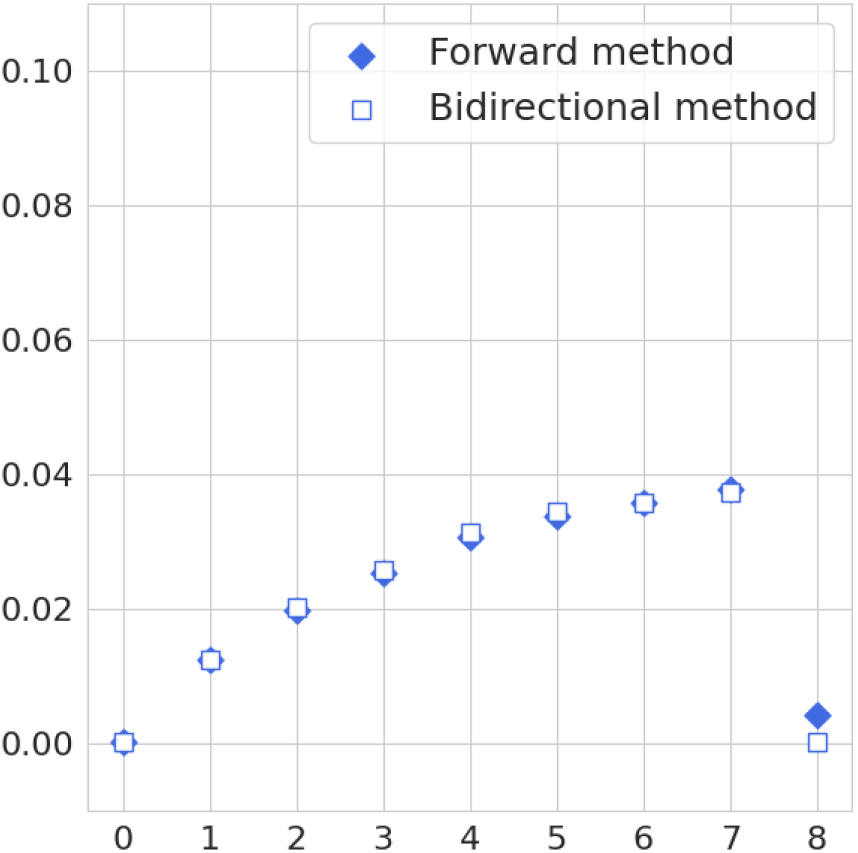
The comparison of the numerical results between forward and bidirectional methods.

This is a simple extension of 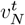 with respect to *t*. The field *v*_ex_(*x, t*) is obtained by applying nearest-neighbor interpolation to 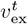.

Hereafter, we present results from numerical experiments for extrapolation using the extended vector field. We performed numerical experiments on two different datasets. The first is composed of the images from No. 0 to No. 10 of HL-60 used in the main paper. In this case, we perform the standard computation between No. 0 and No. 9 and infer No. 10 by extrapolation. Namely, we put *N* = 9 in the definition (12). In the second dataset, we used every other image from No. 0 to No. 10. Concretely, we consider *N* = 4 and each time index *i* corresponds to the image of No. 2*i* in (12).

We assess quantitative performance by comparing our method with an extrapolation procedure based solely on optimal transport (OT) theory. However, to the best of our knowledge, there is no canonical way to perform extrapolation in the optimal transport theory. In this paper, we utilize “*α*-shape” (Edelsbrunner et al. 2003) of the point clouds for conducting the extrapolation within the OT framework.

Here, we provide a brief explanation of the *α*-shape. For a mathematically rigorous and detailed treatment, see (Edelsbrunner et al. 2003). An *α*-shape is the straight-line graph associated with the given data points, and we take its interior as the cell shape. The computation of *α*-shape requires a choice of a parameter *α* ∈ ℝ. This parameter is linked to the *α*-hull, a generalization of the convex hull for a finite point set. For *α* = 0, the 0-hull coincides with the convex hull of the data points, and the *α*-shape is the straight-line graph whose nodes are incident to the vertices of that convex hull.

For *α <* 0, the *α*-hull is the intersection of the complement sets of all circles with radii (−*α*)^−1^ whose complements contain all data points. The *α*-shape is defined as the graph connecting *α*-neighbors, which are edges connecting nodes on the boundary of the same circle with radius −*α*^−1^.

We computed *α*-shape using the functions in the SciPy library (Virtanen et al. 2020). Note that we must choose the parameter *α* to compute the *α*-shape, and its value alters the resulting cell shape. When *α* = 0, the cell shape coincides with the convex hull of the point cloud. If *α >* 0, the *α*-hull is the intersection of closed discs of radius 1*/α*, and the resulting shape becomes convex. However, empirical cell shapes are generally nonconvex; accordingly, we take *α <* 0. If |*α*| is too small or too large, geometric features may not be captured, since the result approaches the convex hull or the original point cloud, respectively. In the simulation, we set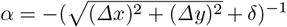, where Δ*x*, Δ*y* are horizontal and vertical spatial step sizes, respectively. The parameter *δ* is a sufficiently small positive real number, which we set to stabilize the extrapolation result. As in the phase-field simulation, the cell shape is represented as an indicator function that equals 1 inside and 0 outside the boundary given by the *α*-shape.

**Figure S4:**
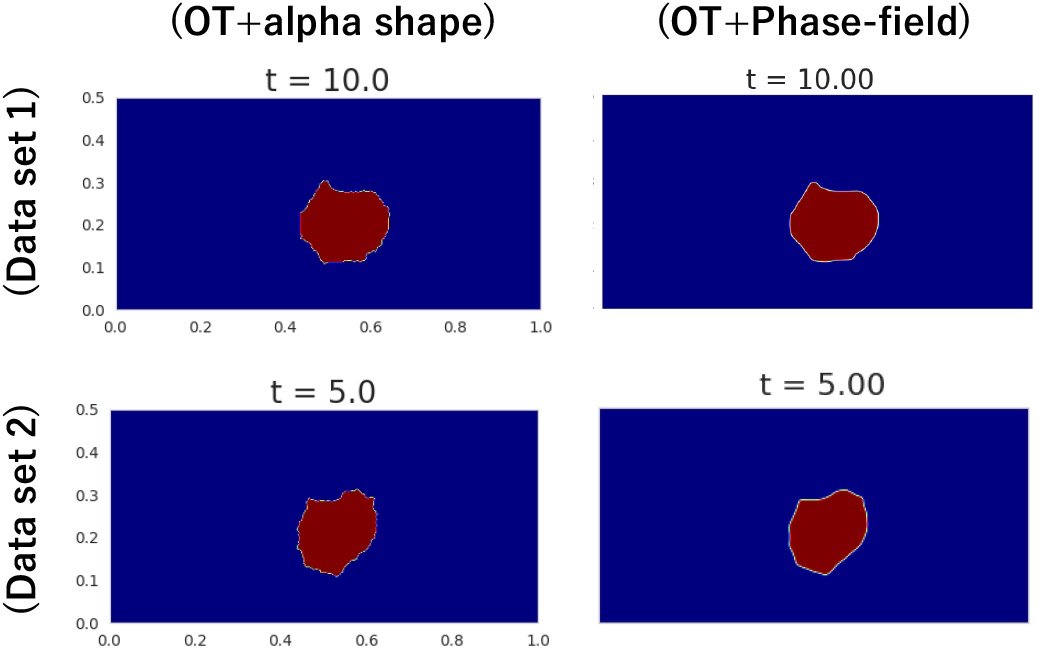
The results of extrapolation by the OT framework and the OT with the phase-field method. The upper and lower rows show the results for datasets 1 and 2, respectively. All pictures show the cell shapes at the final times *t* = 10.0 and *t* = 5.0 for datasets 1 and 2, respectively.

Fig. 4 represents the results of the extrapolation. Under OT with *α*-shape, the cell boundary appears more jagged than with our method. This difference appeared to be induced by the absence of the effect of the interfacial tension in the case of OT with *α*-shape. To compare quantitative performance, we computed the error between the final-time cell shape and the image at time point 10 (the last in the sequence). We used the same error function appearing in the main text. Table 1 reports each value of the errors. In dataset 1, the error from our method is larger than that from OT with *α*-shape. On the other hand, the error obtained by our method is smaller for dataset 2, although the final image is the same in both cases. These observations suggest that the quantitative performance of extrapolation may depend on the dataset.

### G Parameters

Here, we show the parameter settings for the numerical simulation in this paper (Table S2, Table S3).

**Table S1:** Error to image No. 10.

|  | OT | OT + Phase-field |
| --- | --- | --- |
| Dataset 1 | 0.07134 | 0.07715 |
| Dataset 2 | 0.16669 | 0.16547 |

**Table S2:** Parameter setting.

| Parameter | Meaning | Value |
| --- | --- | --- |
| $N_x$ | the number of horizontal grid | the pixel number in horizontal direction of cell shape data |
| $N_y$ | the number of vertical grid | the pixel number in vertical direction of cell shape data |
| $L_x$ | horizontal length | 1.0 |
| $L_y$ | vertical length | $N_x/N_y$ |
| $\Delta x$ | horizontal spatial step size | $L_x/(N_x - 1)$ |
| $\Delta y$ | vertical spatial step size | $L_y/(N_y - 1)$ |
| $\varepsilon$ | order parameter of interfacial width | $1.0 \cdot \Delta x$ |

**Table S3:** Specific parameter setting.

| Simulation | $r$ | Figure |
| --- | --- | --- |
| C.elegans | 0.5 | Figure. 2 in main paper |
| HL-60 | 0.95 | Figure. 2 in main paper |

### H Additional methodological details

#### H.1 Experimental images

Fig. 3(B) employs experimental images of the *C. elegans* embryo obtained by one of the authors through biological experiments. All original *C. elegans* images share the size (*N*_*x*_, *N*_*y*_) = (768, 512), where *N*_*x*_ and *N*_*y*_ denote the numbers of pixels in the horizontal and vertical directions, respectively. Figure 3(C) employs experimental images of HL-60 cells from the supplemental movie in (Imoto et al. 2021). All HL-60 images have the same size (*N*_*x*_, *N*_*y*_) = (238, 292).

#### H.2 Image segmentation

Both the *C. elegans* and HL-60 images were originally grayscale. To extract cell shapes as binary images, image segmentation was applied. The Segment Anything Model (SAM) (Kirillov et al. 2023) was used for segmentation. SAM has been reported to produce high-quality results even on untrained imaging data and may have offered greater flexibility than conventional tools at the time of this study. Images were annotated with reference points indicating the target object, and SAM subsequently performed automatic segmentation based on these points.

For *C. elegans*, reference points were placed near the image center, as the embryo is confined within the eggshell and, at the one-cell stage, the center likely indicates the embryo position. By contrast, HL-60 cells exhibit motility, making manual placement of cell reference points labor-intensive. In this case, the background rather than the cell itself was segmented: a reference point was fixed at a location that the cell appeared not to traverse in the experimental images.

Although SAM generally produced reliable segmentation results in this study, some outputs appeared unrealistic, for example by containing image noise. In such cases, results were corrected manually or refined via morphological transformations using the Python library OpenCV. Specifically, the Closing and Opening operations were used to reduce noise. Closing followed by Opening (CO) was first performed to reduce noise outside the cell shape, and Opening followed by Closing (OC) was then performed to reduce noise inside the cell shape. The resulting cell shape data were regarded as acceptable.

#### H.3 Preprocessing of the cell shape of HL-60

After segmenting the experimental HL-60 images, binary data on a grid of (*N*_*x*_, *N*_*y*_) = (238, 292) were obtained. To promote numerical stability, the interface-width parameter *ε >* 0 was coupled to the grid spacing; accordingly, *ε* was set as *O*(Δ*x*) to simulate interface motion in a stable manner. To allow a smaller *ε*, the binary data were processed as follows. First, the data were resized to (*N*_*x*_, *N*_*y*_) = (292, 292) using zero padding. Second, the data were interpolated to obtain binary data of size (*N*_*x*_, *N*_*y*_) = (1000, 1000). This procedure was not expected to change the aspect ratio of the original cell shape. Finally, during the simulated time interval, the cell appeared to remain in the lower half of the experimental movie; therefore, the upper half of the modified data was clipped to reduce computational cost, yielding binary data of (*N*_*x*_, *N*_*y*_) = (1000, 500), which were used in the numerical simulation in this study.

#### H.4 Numerical algorithm for the phase-field model

We consider the nondimensionalization of (5). The quantities *L*_*x*_ and *L*_*y*_ correspond to the physical length scales of the experimental images. The longer side was rescaled to 1. Namely, if *L*_*y*_ ≤ *L*_*x*_, the normalization *x* → *x/L*_*x*_ and *y* → *y/L*_*x*_ was applied; if *L*_*x*_ *< L*_*y*_, the normalization *x* → *x/L*_*y*_ and *y* → *y/L*_*y*_ was applied. In particular, for the cell shape data used in this study, *L*_*y*_ *< L*_*x*_ was observed to hold, and *L*_*x*_ and *L*_*y*_ are proportional to the grid sizes *N*_*x*_ and *N*_*y*_, respectively. Accordingly, *L*_*x*_ = 1.0 and *L*_*y*_ = *N*_*x*_*/N*_*y*_ were fixed.

For the time scale, the points *t*_0_, *t*_1_, …, *t*_*N*_ were taken to be equally spaced with step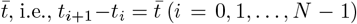. The time variable was then rescaled as 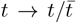, so that the normalized points satisfy *t*_*i*_ = *i* (*i* = 0, 1, …, *N*).

The numerical solution of (5) was computed using an explicit finite-difference method:

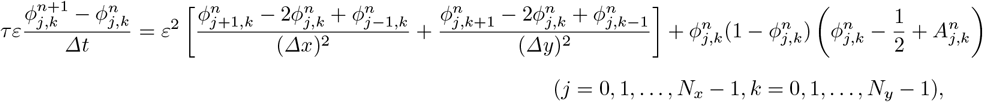

where Δ*t* denotes the time step size, and Δ*x* = *L*_*x*_*/*(*N*_*x*_ − 1), Δ*y* = *L*_*y*_*/*(*N*_*y*_ − 1) denote the spatial step sizes in the *x*- and *y*-directions, respectively. 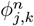 is an approximation of the unknown function *ϕ*, namely 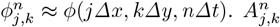 is discretization of *A*_**i**_ in (5) as follows:

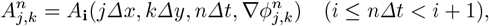

where

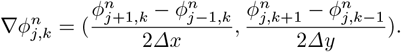

To mitigate potential boundary effects on interface motion, periodic boundary conditions were imposed on (5) and implemented as follows:

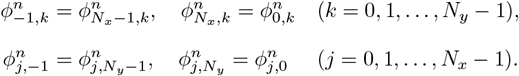

### H.5 Parameter setting

We describe the parameter choices for *ε* and *τ*. The parameter *ε* characterizes the interface width. If *ε* is smaller than the length of the grid, the interface motion may become unstable; therefore, *ε* was set to Δ*x* in this study. The parameter *τ* characterizes the simulation time scale and was set as follows:

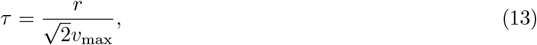

where *r* ∈ (0, 1) is a positive constant and *v*_max_:= max_*t,x,i*_ |*v*_*i*_|. When *τ* satisfies the above, the bistable condition 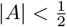 holds due to the definition of *A* and the Cauchy-Schwarz inequality. The quantity *τ*^−1^ is interpreted as the phase-field mobility in the literature of the phase-field method. In this context, (13) suggests that if the maximum of the velocity is large, namely the deformation or movement of the cell is large, the phase-field mobility also needs to be large. Accordingly, the simulation time scale appears to depend on the extent of cell deformation or movement between images.

The constant *r* regulates the maximum phase-field mobility. When *r* is larger, the mobility is smaller. However, if *r* exceeds 1, it may fail to hold the bistable condition. The parameter *r* also appears to moderate the effect of the interfacial tension. Because *A* is chosen to fix the speed of the traveling wave solution in the outer normal direction to ⟨*v*(*x, t*), −∇*ϕ/*∥*ϕ*∥⟩, which is independent of *τ*, the phase-field mobility likely affects only the effect of the interfacial tension.

The value of *r* may be determined by the desired or presumed appropriate cell-shape characteristics. For *C. elegans* (Table S3), *τ* was chosen to be an intermediate value, as the effect of interfacial tension was expected to mitigate the unnatural deformation observed in OT interpolation (Figure S1). On the other hand, for HL-60 (Table S3), *τ* was chosen close to 1 to decrease the effect of interfacial tension. Because the cell is small relative to the computational domain, the cell area may shrink excessively when the interfacial-tension effect is strong.

## Notes

### Competing Interest Statement

The authors have declared no competing interest.

